# Sex-specific remodeling of the human adipose vascular niche in obesity

**DOI:** 10.64898/2026.03.10.710723

**Authors:** Ibrahim AlZaim, Mohamed N. Hassan, Maja Schröter, Bettina Hansen, Henrik Holm Thomsen, Maximilian von Heesen, Lena-Christin Conradi, Martina Rudnicki, Tara L. Haas, Robert A. Fenton, Maria Keller, Matthias Blüher, Niels Jessen, Joanna Kalucka

## Abstract

Obesity remodels the subcutaneous adipose tissue (SAT) vasculature and contributes to cardiometabolic risk, yet potential sex differences in this process remain poorly defined. Here, we integrated single-nucleus transcriptomics and histological analyses of human SAT to reveal pronounced sexual dimorphism within the vascular niche. Obese males exhibit mural cell loss, increased collagen deposition, and inflammatory endothelial activation, including enhanced antigen presentation programs. In contrast, females display preserved mural coverage and increased lipid-handling and redox-adaptive pathways. These coordinated structural and transcriptional differences position the adipose endothelium as a sex-divergent regulator of obesity-associated cardiometabolic vulnerability.

## Introduction

Obesity development and its associated cardiometabolic complications are accompanied by adipose tissue expansion and progressive dysfunction^1,2^. SAT is the primary site of fat storage in humans, and its expansion typically precedes that of visceral adipose tissue (VAT) during the development of obesity^1,3^. Despite this, males with obesity experience a disproportionately higher cardiometabolic risk than females, partly due to a greater propensity for visceral fat accumulation^4^. While this has been partly attributed to differences in fat distribution and inflammatory burden, accumulating evidence suggests that intrinsic biological differences between males and females extend beyond depot anatomy to fundamental aspects of metabolic and vascular regulation^5–8^. These observations have led to a VAT-centric view of obesity-associated cardiometabolic burden. However, more recent evidence suggests that both adipose depots contribute to obesity-associated insulin resistance and cardiometabolic disease risk^9^.

Sexual dimorphism in adipose biology arises from a complex interplay of cellular metabolism, sex hormones, sex chromosome complement, and epigenetic regulation^7,10–13^. These factors influence adipocyte lipid storage capacity, immune cell recruitment, extracellular matrix remodeling, and tissue fibrosis^13,14^. Hallmarks of obesity-associated adipose tissue dysfunction, including adipocyte hypertrophy, vascular rarefaction, immune cell infiltration, and fibrosis, can be observed across depots^2,15,16^. These processes are tightly coordinated within the vascular niche, where endothelial and mural cells integrate metabolic, inflammatory, and fibrotic cues, thereby governing tissue perfusion, immune trafficking, and stromal remodeling^17–20^. Despite this central regulatory role, the vasculature has remained relatively underexplored in the context of sex-specific adipose remodeling.

Although sex differences in adipose tissue distribution and systemic metabolic outcomes are well documented^21^, it remains unclear whether the vascular niche within obese human SAT undergoes sex-specific structural and transcriptional reprogramming. Most prior studies have focused on adipocyte-intrinsic alterations or bulk tissue analyses, limiting resolution of vascular subpopulations and their contribution to sexually dimorphic remodeling. Given the integrative role of the vasculature in coordinating metabolic and inflammatory responses, sex-divergent vascular adaptation could represent a critical yet underappreciated determinant of cardiometabolic risk. We previously generated an integrated single-cell and single-nucleus transcriptomic atlas of the human SAT vascular niche^22^, identifying obesity- and type 2 diabetes–associated vascular phenotypes linked to inflammation and fibrosis. Here, we leverage this resource to determine whether obesity-associated vascular remodeling in human SAT exhibits sexual dimorphism at the cellular, structural, and transcriptional levels.

## Results

### A Single-Cell Atlas Reveals Sex Differences in the Human SAT Vascular Compartment

Recently, we generated a single-cell(sc)/nucleus(sn) transcriptome (sc/sn RNA-seq) atlas of human SAT, comprising nearly 70,000 vascular cells/nuclei derived from 65 donors^22^. More than 60% of the profiled cells originated from female donors, representing approximately 80% of the donor cohort (**Fig. 1a, Supplementary Table 1**). Despite this imbalance, cells from both sexes were detected across all vascular populations. The atlas resolved seven canonical endothelial cell (EC) populations: three venous (V1–V3), two capillary (C1, C2), one arterial (A) and one lymphatic (L), as well as a non-canonical EC population (sub-ECs; S) and two mural cell populations comprising pericytes (P) and vascular smooth muscle cells (SM)^22^. Although both sexes contributed to each vascular subtype, their relative representation differed, as determined by cluster proportions (**Fig. 1b**).

**Fig. 1.**
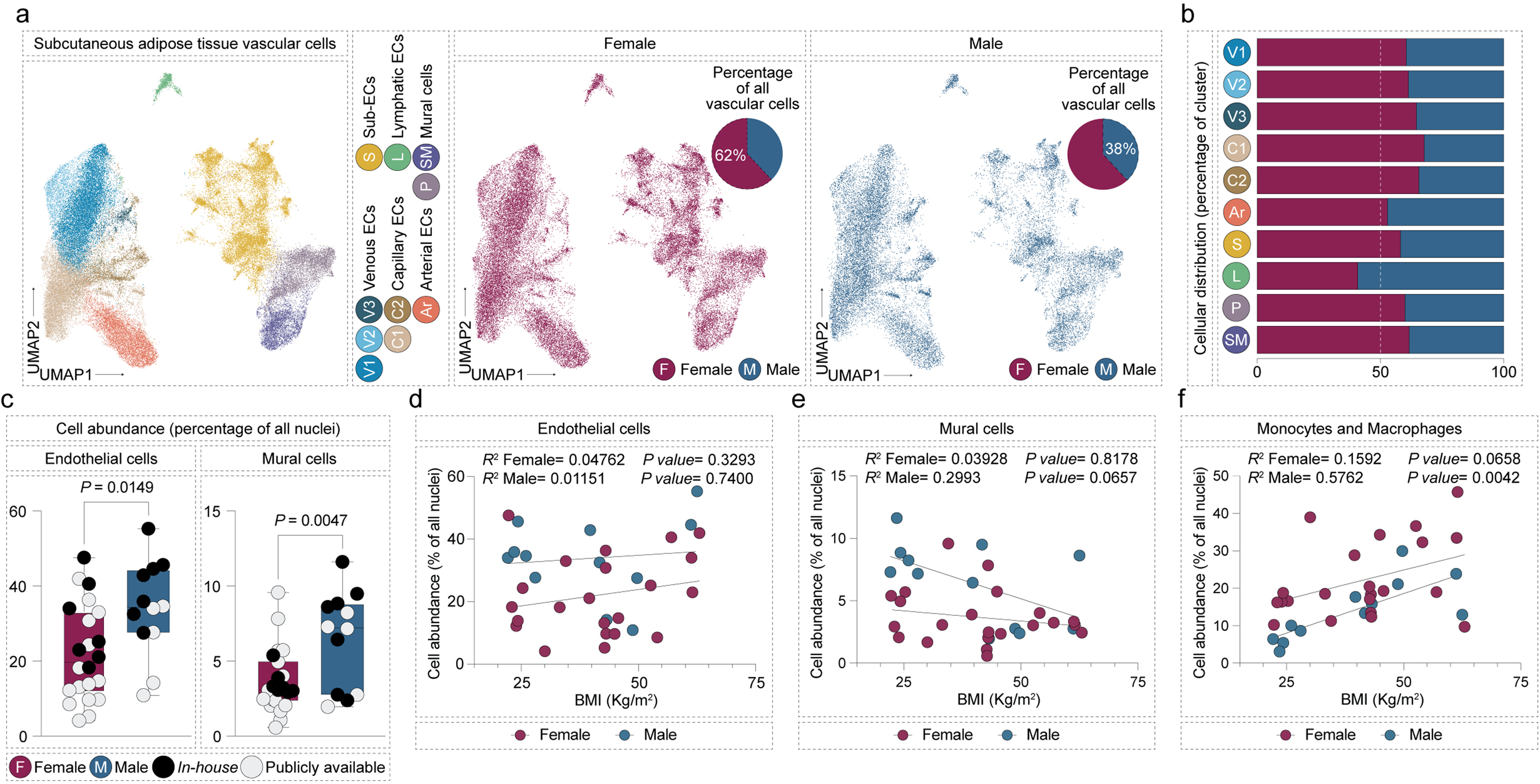
An integrated atlas of human subcutaneous adipose tissue (SAT) identifies sex-specific vascular and immune cell remodeling in obesity. **a,** Uniform manifold approximation and projection (UMAP) depicting the identified cell populations in the SAT vascular cell atlas (*left* panel). UMAPs representing the distribution of the vascular cell populations (Endothelial cells (ECs) and mural cells) within the human SAT vascular cell atlas in females (*central* panel) (51 donors – 62% of all vascular cells) and males (*right* panel) (14 donors – 28% of all vascular cells). Donors included individuals with or without obesity and type 2 diabetes. **b,** Bar graph showing the contribution of either sex to the different vascular cell populations (Venous endothelial cells; V1-3, Capillary endothelial cells; C1,2; Arterial endothelial cells; A, Sub-ECs; S, Lymphatic ECs; L, Pericytes; P, and vascular smooth muscle cells; SM). **c,** Box plots showing the abundance of endothelial and mural cells (as a percentage of all nuclei) from snRNA-seq datasets within the atlas. **d,** Linear regression analysis of the abundance of endothelial cells, mural cells, and Monocytes and Macrophages as a factor of body mass index (BMI). *P value* (c) was calculated using a two-sided unpaired test. For regression analysis (d-f), *P values* were calculated using the two-sided t-distribution. In all cases, *P* < 0.05 was considered statistically significant.

To systematically assess sex-associated differences in cellular composition, we subset the single-nucleus dataset, which enables a comprehensive representation of adipose tissue cellular compartments^23^, and quantified sex-specific changes in cell abundance. Males exhibited a higher abundance of endothelial and mural cell nuclei (as a percentage of all sequenced nuclei per sample) compared to females, independent of metabolic status (**Fig. 1c**). Given prior reports indicating comparable adipocyte size in abdominal SAT between sexes in obesity^24^, these findings suggest that increased vascular cell abundance in males is unlikely to reflect differences in adipocyte hypertrophy. Body mass index (BMI) stratification revealed sex-specific patterns: mural (but not EC) abundance declined with increasing BMI in males, whereas no such association was observed in females (**Fig. 1d,e**). In contrast, monocyte and macrophage abundance was significantly associated with BMI selectively in males (**Fig. 1f**), consistent with the established link between obesity and adipose tissue inflammation^25^. Together, these findings indicate that obesity-associated remodeling of the immune and vascular compartments within SAT is sexually dimorphic.

The higher abundance of endothelial and mural cell nuclei observed in males (independent of the metabolic state) was independently validated by immunohistochemical analysis (**Fig. 2a,c**). Notably, adipocyte size within abdominal SAT has been reported to be comparable between sexes in obesity^24^, suggesting that the higher vascularization observed in males is independent of adipocyte size. Consistent with vascular rarefaction in obesity^2^, overall vessel density was lower in individuals with obesity and with obesity and type 2 diabetes of both sexes (**Fig. 2b**). In contrast, males with obesity exhibited a marked reduction in smooth muscle actin (SMA)-positive mural cell coverage (**Fig. 2d**). This reduction was substantially less pronounced in obese females, revealing sex-specific differences in vascular integrity. Histological quantification showed collagen deposition elevated in obese vs lean SAT, without significant sex differences between obese males and females (**Fig. 2e**), consistent with enhanced fibrotic remodeling of the vascular niche. Taken together, these structural analyses corroborate the presence of sexually dimorphic vascular remodeling in human SAT under obese conditions.

**Fig. 2.**
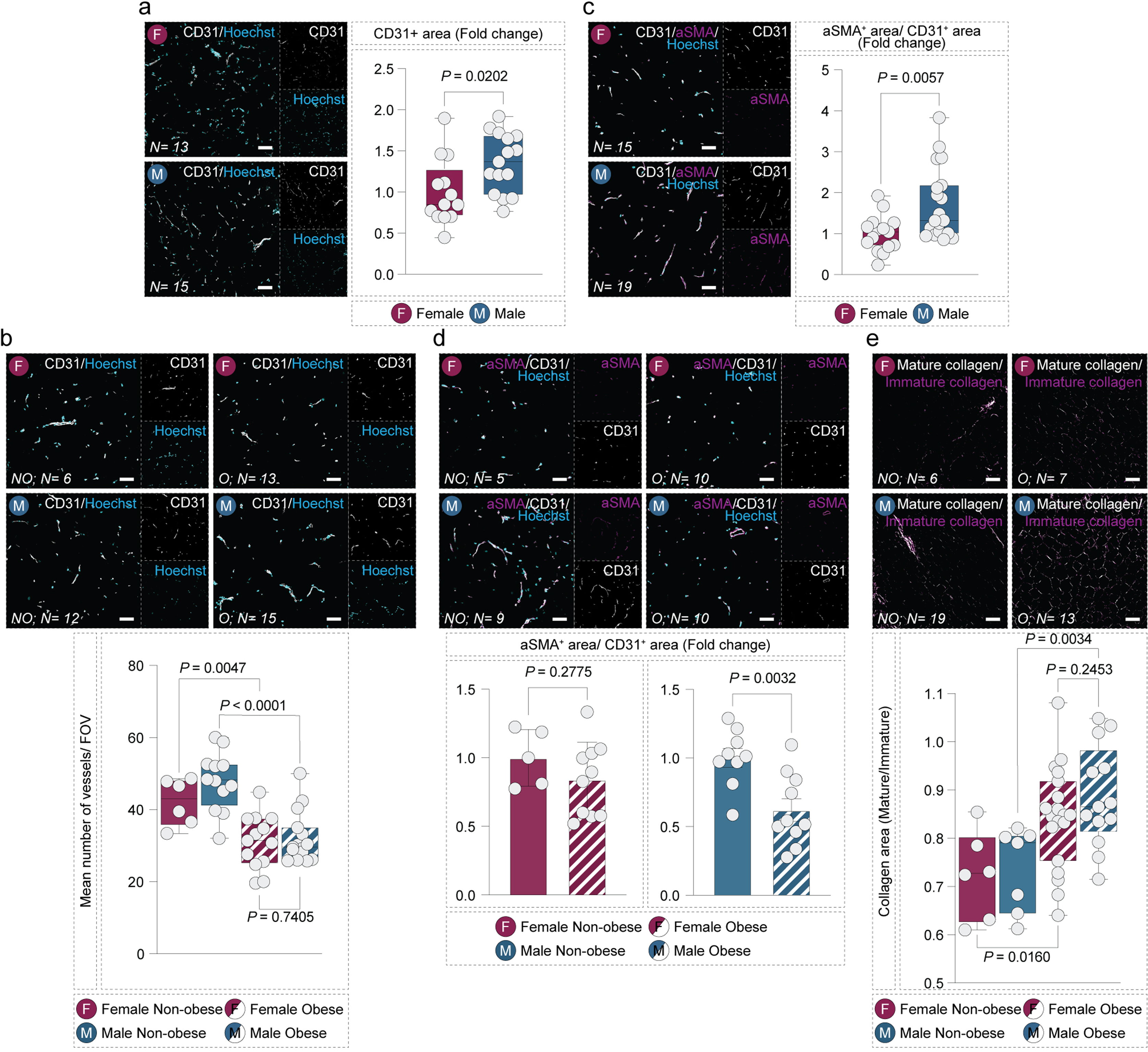
Sex-specific tissue remodeling involves alterations in vascular density, mural cell coverage, and tissue fibrosis. **a,** Representative images and quantification of vascular density, defined as the CD31^+^ area in males and females (Females, *N*=13; Males, *N*=15 distinct donors). **b,** Representative images and quantification of the number of blood vessels per field of view (FOV) in males and females (Female, *N*=6, Female obese, *N*=12; Male non-obese, *N*=13, Male obese, *N*=15 distinct donors). **c,** Representative images and quantification of mural cell coverage, defined as the aSMA^+^/CD31^+^ area (Female, *N*=15; Male, *N*=19). **d,** Representative images and quantification of mural cell coverage, defined as the aSMA^+^/CD31^+^ area (Female non-obese, *N*=5, Female obese, *N*=10; Male non-obese, *N*=9, Male obese, *N*=10). **f,** Representative polarization microscopy images and quantification depicting the ratio of mature to immature collagen as a measure of tissue fibrosis (Female non-obese, *N*=6, Female obese, *N*=19, Male non-obese, *N*=7, Male obese=13). *P values* were calculated using two-sided unpaired t-tests (a, c, and d) or two-way ANOVA (b and e). *P* < 0.05 was considered statistically significant.

### Obesity Drives Sexually Dimorphic Transcriptional Remodeling of the Vascular Niche

To define the transcriptional basis of the remodeling, we performed sex-stratified differential expression analysis (DEA) and gene ontology (GO) analyses across vascular subpopulations using the integrated atlas. Obesity induced both shared and sex-specific transcriptional changes (**Fig. 3a-d, Supplementary Table 3**). Transcriptional remodeling was markedly more extensive in males, with obesity associated with a substantially greater number of differentially expressed genes (DEGs), particularly within capillary and venous ECs (**Fig. 3b, Supplementary Table 3**). In contrast, comparisons between non-obese and obese-diabetic samples revealed only modest additional transcriptional changes in either sex (**Fig. 3a,b**). Collectively, these data indicate that obesity, rather than diabetic status, accounts for the majority of transcriptional variation across vascular subpopulations.

**Fig. 3.**
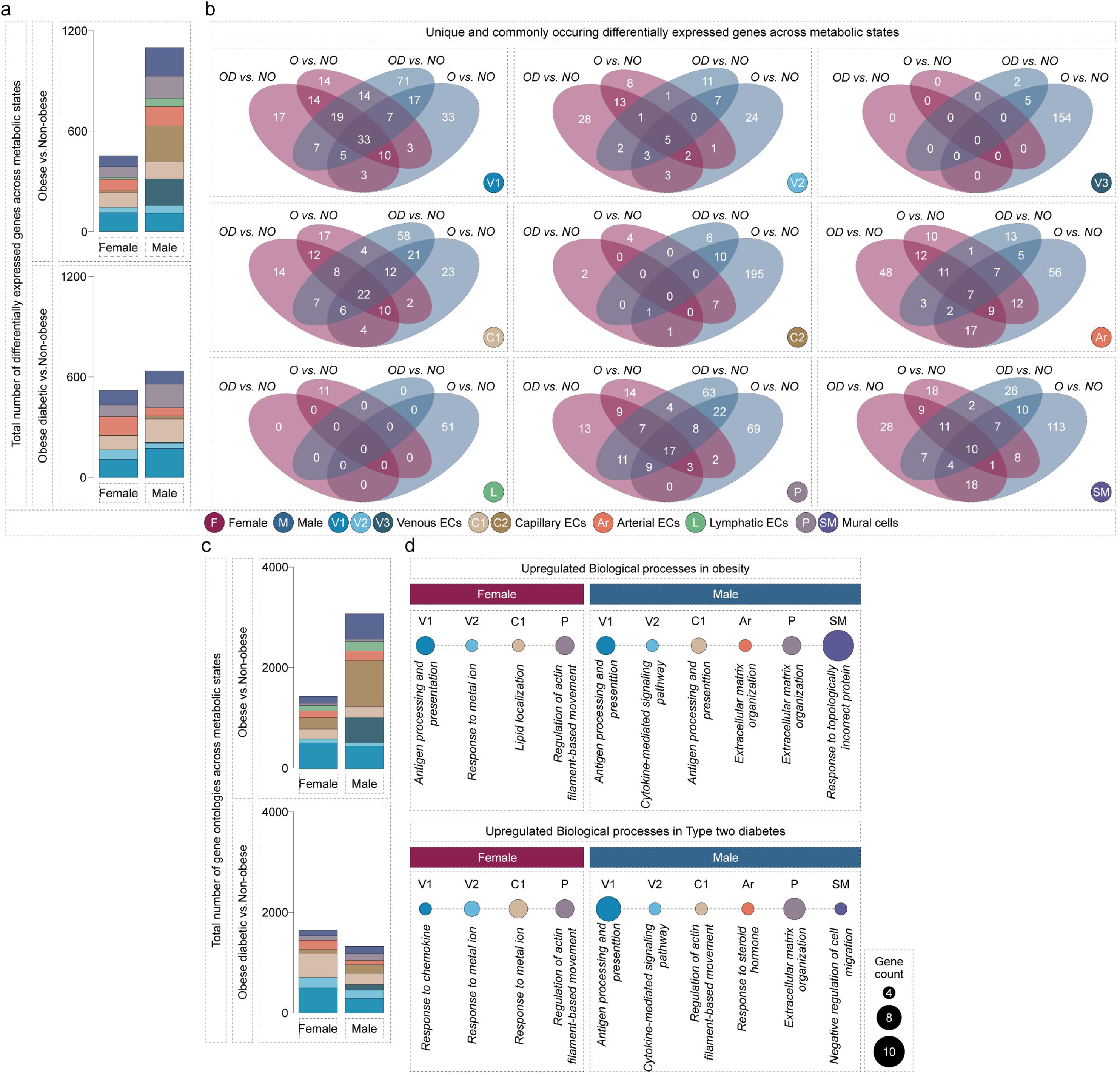
Sex-specific vascular remodeling comprises metabolic state-associated convergent and divergent transcriptomic changes. **a,** Bar graphs showing the total number of identified DEGs in each vascular cell population of either sex across two comparisons (Obese vs. Non-obese and Obese Diabetic vs. Non-obese). **b,** Venn diagrams depicting the number of unique and overlapping DEGs. **c,** Bar graphs showing the total number of identified gene ontology terms in each vascular cell population in females and males. d, Dot plot depicting selected gene ontologies in different vascular cell populations in males and females. Dot size corresponds to gene count.

GO enrichment analyses mirrored this pattern: obese males exhibited stronger pathway enrichment relative to non-obese controls, most prominently within capillary ECs (**Fig. 3c**). Upregulated pathways included antigen processing and presentation, cytokine-mediated signaling, and extracellular matrix organization (**Fig. 3d, Supplementary Table 3**), consistent with the inflammatory and structural changes observed at the histological level (**Fig. 2e**). By contrast, GO enrichment in obese females was comparatively modest and encompassed pro-inflammatory signaling, lipid metabolic processes, and cytoskeletal remodeling pathways. “Response to metal ion” GO-term enrichment signals redox-adaptive endothelial programs under metabolic/inflammatory stress, not metal detoxification, as validated by pathway gene lists (e.g. *MT1A* and *MT2A*). Although several pathways were shared between sexes, the magnitude and functional composition of these programs differed, underscoring pronounced sexual dimorphism. Together, these analyses reveal qualitatively distinct, yet partially overlapping, transcriptional adaptations within the obese vascular niche.

To further dissect the molecular programs underlying these sexually dimorphic signatures, we focused on canonical EC populations. In males with obesity, ECs exhibited marked upregulation of genes encoding *HLA* molecules and *CD74*, implicating these cells in antigen processing and presentation (**Fig. 4a**). Protein-level validation confirmed upregulated HLA and CD74 expression in the vascular endothelium of obese vs lean males, but not females, revealing sex-specific obesity-driven endothelial activation (**Fig. 4b,c**). To assess whether this phenotype was conserved across species, we re-analysed a published snRNA-seq dataset of murine adipose tissue^26^. Male mice fed a HFD displayed increased expression of *Cd74* and genes encoding MHC class II molecules in adipose ECs (**Fig. 4d**). These findings were further recapitulated in reanalyzed sorted adipose ECs from male versus female mice fed a HFD for seven weeks^27^ (**Fig. 4e**), supporting a conserved male-biased induction of antigen presentation pathways under metabolic stress. Notably, the scarcity of HFD single-cell datasets from female mice, limits statistical power for cross-sex comparisons in murine models, a field-wide bottleneck addressed in the Discussion.

**Fig. 4.**
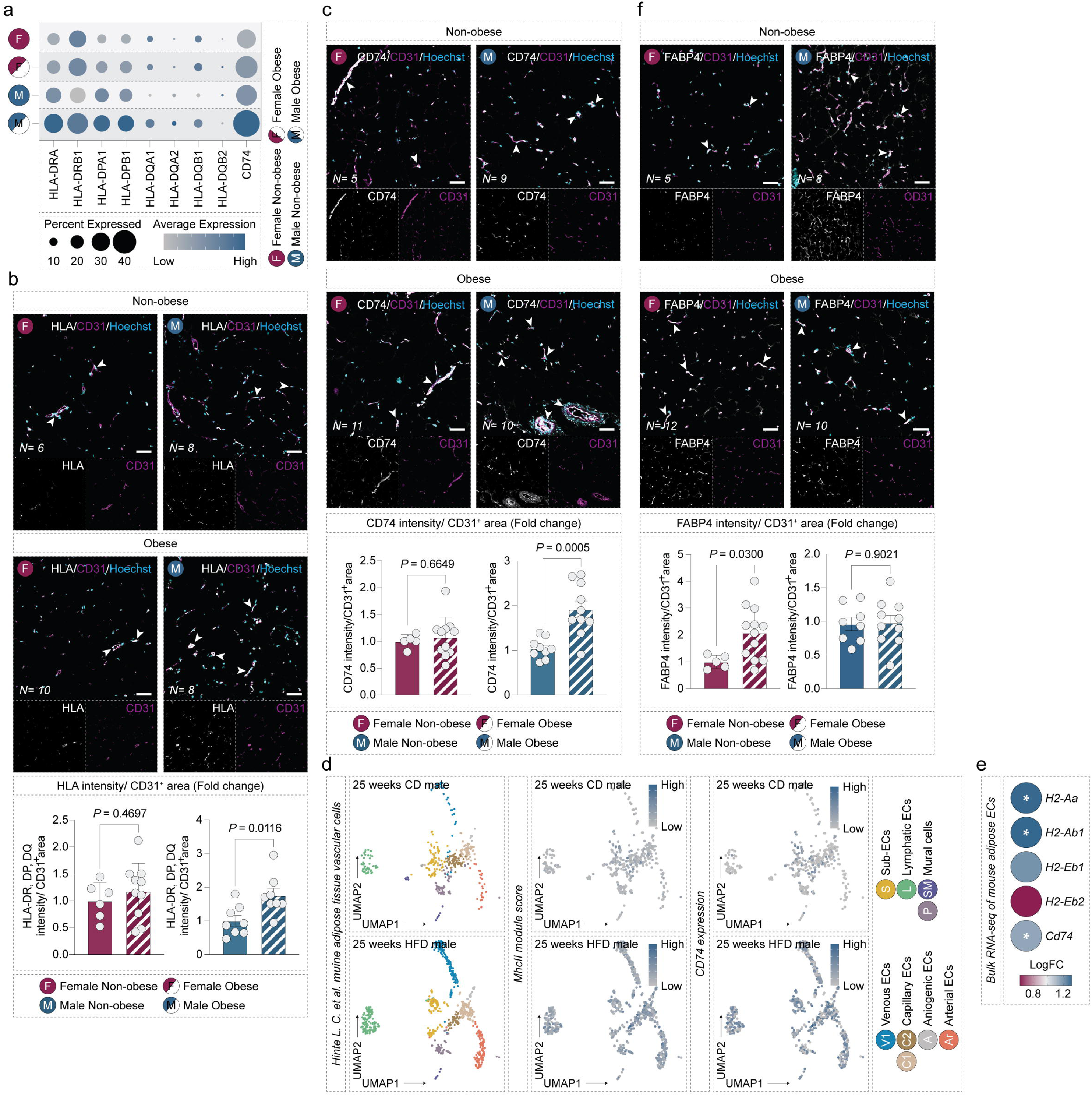
Sex-specific vascular remodeling involves pathways pertinent to antigen presentation and lipid handling. **a,** Dot plot showing the expression of genes encoding HLA class II and CD74 in pooled ECs across sexes and metabolic states (Obese and Obese diabetic cells showed similar trends and were pooled for this analysis). Dot size represents the percentage of cells expressing a certain gene and color reflects average expression level (grey, low; blue, high). **b,** Representative confocal images and quantification depicting the expression of HLA-DR/DP/DQ protein in endothelial cells (CD31^+^ cells) (Female non-obese, *N*=6, Female obese, *N*=10, Male non-obese, *N*=8, Male obese, *N*= 8 distinct donors). **c,** Representative confocal images and quantification depicting the expression of CD74 protein in endothelial cells (CD31^+^ cells) (Female non-obese, *N*=5, Female obese, *N*=11, Male non-obese, *N* =9, Male obese, *N* =10 distinct donors). **d,** UMAP depicting vascular cell populations in adipose tissue of mice fed a control diet (CD) or a high fat diet (HFD) for a duration of 25 weeks (re-analyzed from *Hinte L. C. et al.*). Feature plots showing the *MhcII* score and the expression of *Cd74*. Color reflects average expression level (grey, low; blue, high). **e,** Dot plot showing the enrichment (Log fold change (FC)) of genes encoding MhcII and *Cd74* in sorted adipose tissue endothelial cells in male compared to female mice (re-analyzed from *Rudnicki M. et al.*). **e,** Representative confocal images and quantification depicting the expression of FABP4 protein in endothelial cells (CD31^+^ cells) (Female non-obese, *N*=5, Female obese, *N*=12, Male non-obese, *N*=8, Male obese, *N*= 10 distinct donors). *P values* were calculated using two-sided unpaired t-tests. *P* < 0.05 was considered statistically significant.

In contrast, differential expression and GO analyses revealed enhanced lipid metabolic signatures in ECs of females with obesity. Capillary ECs from females showed increased expression of genes involved in fatty acid handling, including *FABP4*^28^, as well as members of the metallothionein family implicated in protection against oxidative stress (**Fig. 3d**). Upregulation of FABP4 in the vascular endothelium of obese females was confirmed at the protein level (**Fig. 4f**). These data reveal fundamentally distinct endothelial adaptation programs to obesity, characterized by inflammatory activation in males and metabolic–stress adaptation in females.

## Discussion

Obesity remodels the SAT vasculature and contributes to cardiometabolic risk^29^, yet whether these processes diverge between sexes has remained unclear. Here, we demonstrate pronounced sexual dimorphism within the human SAT vascular niche at cellular, structural, and transcriptional levels. Males exhibit a higher baseline abundance of endothelial and mural cells, independent of metabolic status. With obesity, however, males display marked mural cell loss and increased collagen deposition relative to females, consistent with sex-specific microvascular destabilization.

Adipose collagen accumulation is linked to vascular dysfunction and impaired metabolic flexibility, and obesity-associated extracellular matrix (ECM) deposition can localize to the vasculature, altering endothelial behavior^30^. Although direct evidence for sex-specific perivascular ECM remodeling in human SAT is limited, established dimorphism in adipose distribution and inflammatory remodeling supports the plausibility of enhanced collagen deposition in obese male SAT^29–31^. Differences in adipose expansion patterns may further contextualize these findings: females more often exhibit hyperplastic SAT expansion, whereas males tend toward hypertrophic growth^28^. Because collagen deposition by progenitor and fibroblast-like cells constrains adipocyte expandability and increases matrix stiffness^32^, excess ECM accumulation in obese males may compromise microvascular integrity and promote inflammatory activation within the vascular niche.

At the transcriptional level, obesity was associated with broader inflammatory reprogramming in male ECs, including upregulation of antigen presentation pathways such as *HLA class II* genes and *CD74*. In contrast, female ECs preferentially activated lipid-handling and redox-adaptive programs, consistent with enhanced metabolic buffering. These patterns align with HFD studies reporting inflammatory and senescence-associated phenotypes in male ECs and metabolically adaptive signatures in females^27^. Together, our data define a male-biased immunostimulatory endothelial state and a female-biased metabolically resilient phenotype in human obesity.

Collectively, our data position sex as a central determinant of vascular remodeling in obesity. Although the causal contribution of elevated endothelial inflammatory activation to male cardiometabolic risk remains unresolved, most mechanistic obesity studies focus on males, limiting insights into female vascular adaptation. Notably, HFD induces fasting hyperglycemia in males but not females^33^, underscoring divergent metabolic trajectories. Sex-balanced and female-specific models will therefore be essential to interrogate the functional consequences of the dimorphic endothelial programs identified here.

Mechanistically, the vascular divergence observed between sexes likely reflects the integrated influence of endocrine signaling, sex chromosome complement, and sustained differences in metabolic and immune regulation. Estrogen promotes endothelial nitric oxide production and limits vascular inflammation^34,35^, whereas androgens exert context-dependent, potentially pro-inflammatory effects^36^. Differences in X- and Y-linked gene dosage and epigenetic regulation may further preconfigure endothelial and mural cells to respond distinctly to metabolic stress^37^. Although we do not directly dissect these mechanisms, the male-biased antigen presentation signature suggests that sex-specific regulation of interferon and MHC pathways contributes to divergent cardiometabolic trajectories.

Beyond mechanistic insight, these results carry important translational implications. Targeting endothelial antigen presentation or associated cytokine networks may preferentially benefit males, whereas strategies preserving endothelial metabolic resilience may be particularly relevant in females. The limited additional impact of diabetic status on the vascular transcriptional landscape indicates that substantial reprogramming occurs during early weight gain, highlighting a window for intervention. Integrating sex as a biological variable in adipose vascular research will be essential for advancing precision cardiometabolic therapies.

Several limitations warrant consideration. Our analyses are cross-sectional and cannot establish causality. Longitudinal and interventional studies are required to determine whether mural cell loss and antigen-presenting EC expansion precede systemic metabolic deterioration. Functional assays of endothelial-immune interactions and vascular permeability were beyond the scope of this study, and our focus on SAT does not exclude additional sex differences in visceral depots or other organs. Importantly, detailed clinical information regarding menopausal status was not available for the female donors. As ovarian hormone decline profoundly influences metabolic and vascular biology, the ability to stratify samples by pre-versus post-menopausal status would have enabled a more refined interpretation of sex-specific endothelial programs. Mechanistic studies in diet-induced obesity models, particularly incorporating female mice under HFD conditions, will be critical to test causality and conservation of these endothelial programs *in vivo*.

In summary, our findings demonstrate that obesity reprograms the human SAT vasculature in a sexually divergent manner, integrating changes in cellular composition, structural integrity, and endothelial gene expression. In males, obesity is characterized by mural cell loss, fibrosis, and acquisition of an antigen-presenting endothelial phenotype, whereas in females the vascular niche remains structurally preserved and transcriptionally biased toward metabolic adaptation. These data position the adipose endothelium as an active, sex-dependent regulator of cardiometabolic vulnerability and underscore the necessity of incorporating sex as a fundamental biological variable in vascular and metabolic research.

## Methods

### Study cohort and clinical sample acquisition

Human deep SAT samples were obtained from non-obese donors as well as donors with obesity or obesity compounded by type two diabetes undergoing abdominal surgeries at the Clinic for General, Visceral, and Pediatric Surgery of the University Medical Center Göttingen in Göttingen, Germany or the Department of Internal Medicine, Viborg Regional Hospital, Viborg, Denmark. Obesity was defined based on body mass index (BMI) (kg/m^2^): Non-obese donors (BMI < 30) and donors with obesity (BMI > 30). Donors were classified as having type two diabetes based on the presence of a previous history of type two diabetes and antidiabetic medication. All human samples used in the present study were obtained from participants who provided written informed consent according to the University Medical Center Göttingen institutional review board ethical approval (38/4/21) or the scientific ethics committee for region central Jutland approval (1-10-72-312-20).

### Re-analysis of the human SAT vascular cell atlas

An overview of the integrated single cell and snRNA-seq datasets as well as details concerning the generation of the in-house dataset have been previously described^22^. Metrics used for quality control, data integration, and cell type annotation were provided previously^22^. For compositional analysis, only single nucleus data was subset as snRNA-seq faithfully captures the full heterogeneity of the adipose tissue. First, the number of endothelial cell and mural cell nuclei were counted as a percentage of all sequenced nuclei per sample and data was stratified based on sex. Second, scCODA^38^, a Bayesian model for compositional single cell data analysis, was used to validate the sex-associated compositional changes using the snRNA-seq data. Non-obese samples were set as the reference condition and mural cells, being the cluster with the lowest variability across samples and with a low amount of dispersion, were selected as reference. scCODA was ran ten times using the Hamiltonian Monte Carlo sampling method with default parameters. Results were averaged per cell type and average scores were used as a measure of cell type abundance. *FindMarkers* function was used for pairwise comparisons employing a logistic regression test per sex with cohort and chemistry as covariates. The deregulated genes (average log2 fold change above and below 0.5) were then used as input for the enrichGO function from the clusterProfiler package (v4.8.3). All identified genes were used as background for GO enrichment. GO terms of deregulated genes with *P* adjusted values following Bonferroni correction inferior to 0.05 were considered significant. The total number of DEGs and gene ontology terms were visualized using bar plots and the unique and overlapping DEGs across sexes and metabolic states were visualized via Venn diagrams. Selected gene ontologies were visualized in dot plots; scale represents gene count per GO term.

### Re-analysis of snRNA and bulk RNA-sequencing data of murine adipose tissue endothelial cells

Publicly available snRNA-seq data of murine adipose tissue^26^ were preprocessed (500 > *nFeature_RNA* > 4000, *nCount_RNA* < 40000, and percentage of mitochondrial read < 20%). Confounding genes including mitochondrial, ribosomal, and hemoglobin genes as well as *Malat1* and *Neat1* were removed. Outliers were removed based on the number of Unique molecular identifiers and genes detected. Potential doublets were removed using DoubletFinder^39^ (v2.0.3). Data were merged, normalized using *NormalizeData* then multiplied by a factor of 10000. Top 2000 highly variable genes were selected and scaled regressing out the number of UMIs. Dimensionality reduction was performed using principal component analysis (PCA) and projected using Uniform Manifold Approximation and Projection (UMAP). Batch correction was performed using *Harmony*^40^ (v0.1.1), specifying samples as a covariate. Clustering was performed on 43 *Harmony*-corrected components and cells were clustered at a resolution of 0.1. Clusters were annotated based on their expression of canonical marker genes^41^ and vascular cells (endothelial cells, lymphatic endothelial cells, and mural cells) were subset for downstream analysis. Vascular cells were processed as described above and clustered at a resolution of 0.7. Clusters were annotated based on previously reported marker genes and clusters with no noteworthy differences were collapsed. The *AddModuleScore* function was used to calculate the *MhcII* score using genes encoding MhcII (*H2-Aa*, *H2-Ab1*, *H2-Ea*, *H2-Eb1*, and *H2-Eb2*). Data corresponding to mice fed a control or a high fat diet are presented. Bulk RNA-seq data of male and female murine adipose tissue ECs (sorted from HFD mice) were provided by T. Haas^33^. LogFC values (male vs. female) were used for validation against our scRNA-seq data.

### Histology and Immunohistochemistry

Paraffin embedded human adipose tissue blocks were sectioned using an automated microtome HM 355S (Epredia) into 5 μm sections. For collagen detection, picrosirius red staining was performed. Sections were deparaffinized overnight in xylene and rehydrated in graded ethanol. Staining was performed using a commercial kit (24901-500, Polysciences), following which, sections were dehydrated and mounted. Sections were imaged (4-5 random fields of view (FOV) per section) using a brightfield microscope equipped for polarization microscopy with a crossed polarizer, an AxioCam506 detector (6Mpx color camera, 4.54μm/pixel), a manual rotating stage, and an N-Achroplan 10x/ 0.25 Pol objective. Polarized light allows the visualization of collagen fiber structure and maturity, discriminating between mature and immature collagen based on fiber thickness packing density, and degree of crosslinking. Mature collagen corresponds to collagen I-rich fibers and reflects established fibrosis while immature collagen corresponds to collagen III-rich fibers and reflects early pathological ECM expansion. For immunohistochemistry, sections were deparaffinized in Neoclear and rehydrated in graded ethanol then subjected to antigen retrieval in TEG buffer (0,01M Tris (77-86-1, Sigma Aldrich), 0,5mM EGTA (67-42-5, Merck) in double distilled water, pH 9) in a steamer for 60 minutes. Sections were allowed to cool to room temperature after which sections were washed and incubated with protein blocking buffer (3% BSA, 0,1% Triton-X100 in PBS) for one hour at room temperature. Sections were then incubated with primary antibodies diluted in protein blocking buffer (3% BSA, 0,1% Triton-X100 in PBS) at 4 degrees overnight, washed with PBS, and incubated with 1:1000 secondary antibodies diluted in protein blocking buffer for 2 hours at room temperature. Tissues were then washed with PBS and counterstained with Hoechst (1:5000) prior to mounting. The following primary antibodies were used: 1:800 mouse anti-CD31 (3528, Cell Signaling Technology), 1:100 rabbit anti-CD31 (NB100-2284, Novus Biologicals), 1:200 mouse anti-alpha smooth muscle actin (M085129, Agilent), 1:1000 HLA-DR, DP, DQ (555556, BD Pharmigen), 1:1000 rabbit anti-CD74 (77274, Cell Signaling Technology), 1:400 and rabbit anti-FABP4 (710189, Invitrogen). Secondary antibodies used were donkey anti-mouse AF568 (A10037, Invitrogen), donkey anti-rabbit Alexa Fluor 555 (A31572, Invitrogen), donkey anti-mouse Alexa Fluor 555 (A-31570, Invitrogen), donkey anti-mouse AF488 (A21202, Invitrogen), and donkey anti-rabbit Alexa Fluor 488 (A-21206, Invitrogen). Z-stack images (4-5 random FOV per section) were acquired using a LSM 900 inverted confocal microscope equipped with an Airyscan 2 detector array, 4 lasers (405nm, 488nm, 561nm, and 640nm), and a Pln-Apo 10x/045 objective. Images were exported as maximum intensity projections using the Zeiss software. Image analysis was performed using Fiji (v.2.14.0) as described below.

### Image analysis

Vascular density was quantified as the CD31^+^ area and mural cell coverage as aSMA^+^ area/CD31^+^ area per image. First, a threshold-based mask was applied to the relevant marker, generating ROIs of which the area was measured. The number of vascular structures per FOV were also manually counted and presented as a mean number of vessels/ FOV. Alternatively, the intensity of proteins of interest within the vascular compartment were quantified in vascular ROIs in which the vessel-based ROIs were applied to the channel containing the relevant marker of interest and the mean gray value (MGV) of the signal was calculated within the ROIs after the channels were split. Ratios were calculated by normalizing the average MGV of the marker of interest to the average area of the relevant vessel marker. Mature and immature collagen were distinguished leveraging the birefringent properties of collagen. Image acquisition settings (exposure, gain, and white balance) were kept constant across samples to allow comparative analysis and data was presented as the ratio of mature to immature collagen.

### Statistics

Statistical analysis was performed using GraphPad Prism v.10.2.2 and data are represented as mean ± s.e.m. Statistical significance between two groups was calculated using an unpaired two-tailed t-test with Welch correction. Linear regression analysis was used to assess the relationship between nuclei abundance and BMI. *P* values inferior to 0.05 were considered significant. Statistical tests used for all computational analyses were previously described^22^. Gene ontology analysis was performed using over-representation analysis based on hypergeometric distribution. For computational analyses, *P* values were corrected for multiple testing using the Benjamini-Hochberg method and were considered significant if inferior to 0.05.

